# Dissecting the strain and sex specific connectome signatures of unanesthetized C57BL/6J and DBA/2J mice using magnetic resonance imaging

**DOI:** 10.1101/2025.09.23.678044

**Authors:** Tanzil M. Arefin, Hayreddin Said Unsal, Thomas Neuberger, Nanyin Zhang, Helen M. Kamens

## Abstract

Mouse models are an essential tool for understanding behavior and disease states in neuroscience research. While genetic and sex-specific effects have been reported in many neurodegenerative and psychiatric illnesses, these factors may also alter baseline neuroanatomical features of mice. This raises the question of whether the observed changes are related to the disease being studied (i.e., pathological differences) or if there are baseline strain or sex differences that may potentially predispose animals to different responses. Over the past decade, tremendous effort has been made in mapping neural architecture at various scales; however, the complex relationships including identifying genetic and sex-specific differences in brain structure and function remain understudied. To bridge this gap, we used C57BL/6J and DBA/2J mice, two of the most widely used inbred mouse strains in neuroscience research, to investigate strain and sex-specific features of the brain connectome in awake animals using magnetic resonance imaging (MRI). By combining resting-state fMRI and diffusion MRI, we found that the motor, sensory, limbic, and salience networks exhibit significant differences in both functional and structural domains between C57BL/6J and DBA/2J mice. Further, functional and structural properties of the brain were significantly correlated in both strains. Our results underscore the importance of considering these baseline differences when interpreting the brain-behavior interactions in mouse models of human disorders.

## Introduction

During development, newly born neurons migrate, differentiate, and form distinct anatomical units and sophisticated functional networks via axonal connections. Emerging studies suggest that the molecular mechanisms govern the functioning of the brain through which specific behavior or pathology is expressed. Importantly though, these mechanisms can be altered by many factors, including the sex of the animal or the genetic makeup [1–7]. Therefore, examining the influence of such factors on neuro-architecture is important not only for understanding normal brain function but also for understanding the neural mechanisms that lead to behavioral or disease states.

Inbred mice have been widely used to study the genetic and molecular mechanisms underlying complex diseases due to the accessibility of mouse genomic sequence and a large number of publicly available resources detailing physiological and behavioral changes [8, 9]. C57BL/6J (C57) and DBA/2J (DBA) mice are two of the most investigated strains in neuroscience research. C57 mice are an ideal testbed for examining the molecular mechanisms underlying diseases since it is the standard genetic background on which most gene knockouts have been developed, and its genome was the first to be sequenced [10, 11]. DBA, on the other hand, is often used as a contrast to C57 mice because they have divergent phenotypes for many traits. Numerous studies have provided evidence of divergent phenotypes between C57 and DBA male and female mice for outcomes such as neurodegenerative and psychiatric illnesses, including addiction [12–17]. This raises the question of whether the observed changes are related to the disease being studied (i.e., pathological differences) or if there are baseline strain or sex differences that may potentially predispose animals to different responses. Hence, unveiling genetic and sex contributions to baseline brain structure and function is important.

Electrophysiology, tissue histology, and currently two-photon tomography have been considered the gold standard for studying neural activity and the microstructure of the brain. However, these techniques typically examine a specific region or pathway in a single brain and do not allow us to examine multiple circuits within the same brain. Therefore, a quantitative evaluation of the neuronal organization of multiple strains or between male and female animals remains a challenge. In contrast, magnetic resonance (MR) based imaging techniques, such as resting-state functional MRI (rsfMRI) and diffusion MRI (dMRI), allow for non-invasive mapping the function and structure of multiple circuits across the entire brain. Furthermore, by utilizing computational tools, we can combine rsfMRI and dMRI to non-invasively elucidate the microstructural fingerprints underlying the functional networks within the same animal. These techniques are ideal for detailing the complex structural-functional relationships in the rodent brain and the factors that modify these relationships [6, 18–22].

To date, several neuroimaging modalities have been used to map the strain- and sex-specific patterns of the structural and functional connectome of the mouse. For example, using rsfMRI and dMRI, strain-specific structural and functional connectivity have been reported in the C57BL/6N and BALB/cJ mice [23]. The heritability of the brain microscopic structures and parameters were estimated from a diverse cohort of mice (C57BL/6J, DBA/2J, CAST/EiJ, and BTBR T + *Itpr3^tf^*/J) using ultra high-resolution dMRI indicating a genetic contribution to these traits [24]. Furthermore, cross-species convergence of sex differences in mouse and human brain anatomy has been reported using structural magnetic resonance imaging (MRI) [25]. However, if there are differences in the brain connectome specific to strain and sex in C57 and DBA mice has not been examined. To bridge this gap, we combined rsfMRI and dMRI to investigate genetic and sex-specific signatures of the brain connectome in C57BL/6J and DBA/2J male and female mice.

A broader aim of this study was to establish an imaging framework for the quantitative measurement of genetic and sex-specific signatures in the brain connectome, thereby understanding the causative link between structure and function. Combining non-invasive rsfMRI and dMRI techniques, we addressed three questions. First, what are the differences in the spontaneous neural activity and the structural integrity of the brain between C57 and DBA mice and between male and female mice? Second, how do strain and sex affect functional and structural connectivity? Finally, is there a relationship between structural and functional properties of the brain?

To answer these questions, we first, measured spontaneous neural activity of the brain by examining the amplitude of low frequency fluctuations (ALFF) of the blood oxygen level-dependent (BOLD) signal across the whole brain. We then measured the structural fractional anisotropy (FA), quantifying the degree of water diffusion in the brain. Subsequently, we quantified the significant effects of strain, sex and the interaction of these factors on FA and ALFF. This hypothesis free approach allowed us to detect the brain regions that show significant differences when considering these factors. Second, using the brain regions identified with the hypothesis free approach as nodes, we computed node-to-node functional and structural connectivity (FC and SC). Here, we again examined the effects of strain and sex on these networks. Finally, we identified the structural-functional relationship of brain regional properties (i.e., FA and ALFF) as well as in the node-to-node communications (i.e., SC and FA). Our findings highlight the significant effects of strain and sex on the brain connectome, which should be considered as researchers use these models to understand the neural basis of behavioral and disease states.

## Methods

### Animals

All experiments were approved by and conducted in accordance with guidelines from the Pennsylvania State University Institutional Animal Care and Use Committee (IACUC). C57BL/6J (male: N = 10, female: N = 10) and DBA/2J (male: N = 10, female: N = 10) mice were obtained from The Jackson Laboratory, Bar Harbor, ME, USA. Animals were group-housed (2-4/cage) in Plexiglas cages. The animal facility was maintained 21-22°C and 55-65% humidity under a 12-hour light/dark cycle (lights on at 7:00 AM) with water and chow available *ad libitum*.

### Experimental design

#### Animal habituation

Fig. 1a demonstrates the experimental paradigm used for this study. All animals underwent a 7-day acclimation procedure followed by imaging sessions in the awake (unanesthetized) state as described in our earlier studies [26–28]. In brief, each mouse was habituated to the scanning environment using a custom-designed restrainer while MRI noises were played to minimize stress and motion during the imaging session. The duration of training gradually increased from 15 minutes (min) on day 1 (D1) to 30 min on D2, 45 min on D3, and 60 min during D4 – D7. Functional and anatomical images were acquired 48 hours after the last habituation day (D9). This gradual habituation was previously shown to reduce hypothalamic-pituitary-adrenal (HPA axis activation [29].

**Fig. 1:**
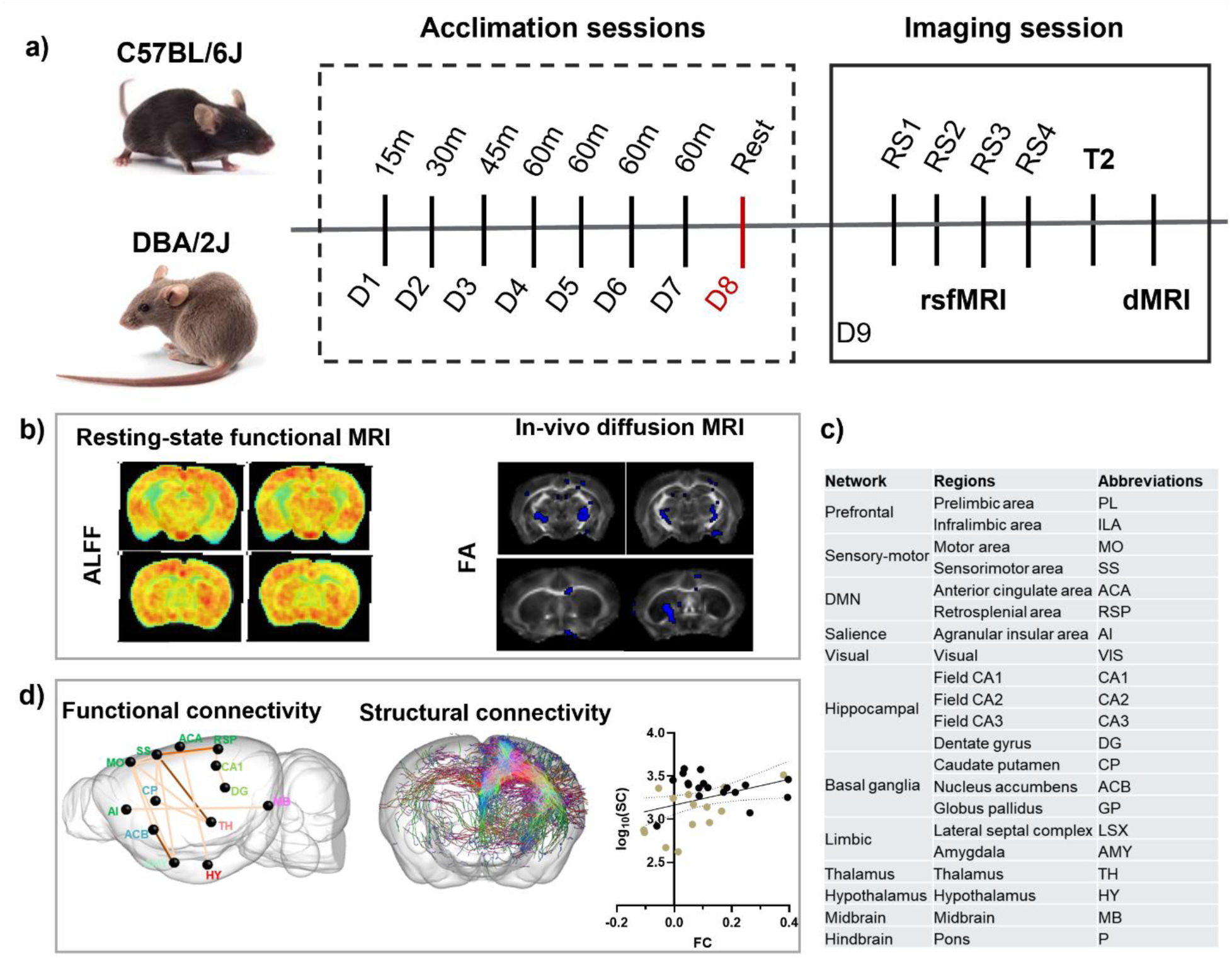
Study overview: a) Experimental timeline: Male and female C57BL/6J and DBA/2J mice were used in this study (N = 10 /group). Each animal underwent habituation session for 7 consecutive days, starting 15 minutes (m) on day 1 (D1), 30m on D2, 45m on D3, followed by 60m for next 4 days (D4 – D7). 48 hours after the last habituation session, resting state functional Magnetic Resonance Imaging (rsfMRI) data were acquired from 4 consecutive scans followed by a structural MRI (T₂) and diffusion MRI scan. b) Resting spontaneous neuronal activity was quantified by amplitude of low frequency fluctuation (ALFF) and microstructural integrity by fractional anisotropy (FA). c) Strain and sex dependent effects on ALFF and FA were observed in 21 regions. Finally in d) strain- and sex-dependent effects on functional and structural connectivity were assessed.

#### MRI data acquisition

Animals were briefly exposed to isoflurane for placement into the restrainer; however, isoflurane was discontinued immediately after head restraining. The animals were then inserted into the scanner. Prior to the imaging sessions, positioning of the animal inside the bore and the typical scanner adjustments (frequency, shim, Bo map, and reference power) took about 15 - 20 minutes. Therefore, animals were fully alert by the time the rsfMRI session started. MRI data were acquired with an in-house built single-channel loop coil (20 mm diameter) for MR signal transmission and reception using a 7T Bruker 70/30 BioSpec system (ParaVision 6.0.1, Bruker, Billerica, MA) at the Penn State High-Field MRI Facility. rsfMRI data was recorded in 4 consecutive sessions, followed by an anatomical multi-slice T_2_-weighted scan utilizing the turbo rapid acquisition with relaxation enhancement (Turbo-RARE) method for each mouse. Subsequently, diffusion MRI (dMRI) data was acquired under isoflurane (∼1.5 vol%) on respiration triggering. Isoflurane was used during diffusion imaging to avoid movement artifacts. Physiological conditions (temperature and respiration) were continuously monitored during the imaging session using a small animal monitoring system (SAI, Inc., Stony Brook, NY).

#### Resting-state functional MRI (rsfMRI)

Awake mouse rsfMRI data were acquired using a T_2_*-weighted single-shot multi-slice gradient-echo echo planar imaging (GE-EPI) sequence [18, 21] (echo time (TE) = 15 ms, repetition time (TR) = 1500 ms). The entire mouse brain was covered using 16 axial slices of 0.75 mm thickness acquired in an interlaced manner, with a field of view (FOV) of 16 × 16 mm^2^, and an acquisition matrix of 64 × 64, which resulted in an in-plane resolution of 250 × 250 µm^2^. 400 image volumes were acquired for each ∼10min run.

### Anatomical imaging

Following rsfMRI, morphological images were acquired using a multi-slice T_2_-weighted Turbo-RARE sequence [30] (TE/TR = 8 ms/1500 ms, six averages, RARE factor of 6). Like rsfMRI, the entire mouse brain was covered using 16 slices of 0.75 mm thickness with a FOV of 16 × 16 mm^2^, and acquisition matrix of 128 × 128, resulting in a planar resolution of 125 × 125 µm^2^.

### In-vivo diffusion MRI (dMRI)

multi-shell dMRI data were acquired using a four-shot multi-slice diffusion tensor imaging-EPI (DTI-EPI) sequence [18, 21] with *b* factors of 800 s/mm^2^ and 2000 s/mm^2^ in 60 noncollinear diffusion gradient directions per b-value.

16 axial slices of 0.75 mm thickness were acquired at a spatial resolution of 125 × 125 μm^2^ with an FOV of 16 × 16 mm^2^ and an acquisition matrix of 128 × 128, and the following diffusion parameters: TE/TR = 25 ms/6000 ms, diffusion gradient separation (Δ) = 10 ms, diffusion gradient duration (*δ*) = 3.5 ms. During dMRI, we used same orientation and geometry of the slices as the rsfMRI to cover similar anatomical regions across scans.

### rsfMRI data pre-processing

rsfMRI data were reconstructed and preprocessed (Fig. 1b) using in-house developed MATLAB R2023a (www.mathworks.com) scripts [31]. In brief, the following steps were applied for each animal:

#### Despiking

The first 10 volumes of each raw rsfMRI dataset were discarded to ensure the steady state of magnetization. Volumes with excessive motion of the remaining scans were discarded by computing framewise displacement (FD) using the method described in [31, 32]. Volumes with FD > 0.2 mm, along with their adjacent volumes, were removed. Any subject showing more than 10% image volumes with >0.2mm FD was flagged as ‘failed to adapt to the restrainer or scanner environment’, i.e., an outlier. We detected two outliers (C57 – male: N=1, and DBA – male: N = 1) under this condition; however, replacement animals were used to match the number of subjects per condition.

#### Image coregistration

Functional and anatomical data were coregistered using the image coregistration pipeline described in our previous study [18, 19, 33, 34]. First, rsfMRI data were mapped to the respective T_2_ image volume using a normalized mutual information approach, a 4th degree B-Spline interpolation, and a 6-parameter rigid body transformation (3 parameters for translation and 3 parameters for rotation). rsfMRI and the respective T_2_ data were then transferred into a template space (150 µm³ isotropic resolution, 54 axial slices) aligned with the mouse atlas [19] using automated image registration (AIR) followed by large deformation diffeomorphic metric mapping (LDDMM) transformation implemented in DiffeoMap (www.mristudio.org).

#### Motion correction

Motion related artifacts were corrected by realigning rsfMRI volumes to the first volume using a least-square approach with a 6-parameter rigid body spatial transformation as implemented in statistical parametric mapping (SPM) [18, 31].

#### Nuisance regression

Brain white matter (WM) and cerebral spinal fluid (CSF) masks from the atlas were mapped onto each subject’s rsfMRI images. Nuisance signals from the brain WM and CSF, as well as motion parameters, were regressed out voxel-wise from the BOLD time series [31].

#### Smoothing and filtering

Finally, rsfMRI images were spatially smoothed (Gaussian kernel, FWHM = 1 mm) and temporally filtered by a 0.01–0.1 Hz bandpass filter [18, 31].

### dMRI data pre-processing

dMRI data were pre-processed as described in our earlier studies[19, 33]. Briefly, non-brain tissue signals were discarded manually from the dMRI data using AMIRA (ThermoFisher Scientific, https://www.thermofisher.com) and corrected for ghosting artifacts likely caused by frequency drifts using rigid alignment implemented in DTIStudio (32).

### Anatomical labeling of the MRI data

Both the rsfMRI and dMRI data were coregistered into high-resolution diffusion MRI based mouse brain atlas [19] using rigid transformation followed by intensity-based affine and LDDMM. Detailed mouse brain structural labels containing 70 brain regions were transferred to each brain individually, as described previously [19, 33]. Subject-specific brain regions were used to compute the brain structural and functional properties of each mouse individually.

### MRI data post-processing

Strain- and sex-specific signatures of the brain connectome were assessed in two steps. First, we performed a regional analysis. We used two imaging metrics – amplitude of low frequency fluctuation (ALFF) from rsfMRI and fractional anisotropy (FA) from dMRI respectively, to quantify functional and structural parameters.

*<u>ALFF:</u>* We used ALFF to quantify the spontaneous neural activity in the brain during rest by measuring the fluctuations of BOLD signal in the range of 0.01-0.1 Hz [35]. First, the time course of each voxel was converted to the frequency domain using a fast Fourier transformation (FFT). Then, the ALFF was measured by computing the square root of the power spectrum and averaging throughout the bandpass (0.01–0.1 Hz) at each voxel. The ALFF of each voxel was divided by the global ALFF value for standardization purposes. Finally, the group averaged ALFF maps were generated individually [4] (Fig. 1b).

*<u>FA:</u>* We computed FA to examine the degree of overall directionality of water diffusion that reflects the white matter (WM) integrity in the brain (Fig. 1b). First, the six elements of diffusion tensor of the brain were determined by log-linear fitting via DTIStudio [36]. Next, three eigenvalues (λ_1_, λ_2_, λ_3_) and corresponding eigenvectors (v_1_, v_2_, v_3_) were computed by diagonalizing the tensor, and FA was then calculated from the eigenvalues [37, 38] as follows:

√3/2 × [√ ((λ_1_ − λ)^2^ + (λ_2_ − λ)^2^ + (λ_3_ − λ)^2^) / √ (λ_1_^2^ + λ_2_^2^ + λ_3_^2^)], where λ_n_ = the eigenvalues describing the diffusion tensor, and λ is the mean diffusivity ((λ_1_ + λ_2_ + λ_3_)/3). Higher FA values indicate stronger WM integrity and vice versa.

#### Selection of nodes

Examining regional FA and ALFF, we identified 21 brain regions that exhibited significant effects (2 × 2 ANOVA, α = 0.05, FDR corrected) of strain or sex, which were used for further analysis (Fig. 1c).

#### Resting state functional connectivity (rsFC)

We used the 21 bilateral nodes (table 1) to compute the rsFC between each pair of nodes, i.e., node-to-node rsFC for each subject. The time course of each node was obtained by averaging the time courses of all voxels within the node. Node-to-node rsFC was calculated using Pearson correlation coefficient between their time courses, which was then converted to a z score using Fisher’s r-to-z transform (Fig. 1d).

**Table 1:** List of the regions showing significant changes between strains or sexes based on ALFF or FA. DMN = default mode network.

| Network | Regions | Abbreviations |
| --- | --- | --- |
| Prefrontal | Prelimbic area | PL |
|  | Infralimbic area | ILA |
| Sensory-motor | Motor area | MO |
|  | Sensorimotor area | SS |
| DMN | Anterior cingulate area | ACA |
|  | Retrosplenial area | RSP |
| Salience | Agranular insular area | AI |
| Visual | Visual | VIS |
| Hippocampal | Field CA1 | CA1 |
|  | Field CA2 | CA2 |
|  | Field CA3 | CA3 |
|  | Dentate gyrus | DG |
| Basal ganglia | Caudate putamen | CP |
|  | Globus pallidus | GP |
| Limbic | Nucleus accumbens | ACB |
|  | Lateral septal complex | LSX |
|  | Amygdala | AMY |
|  | Thalamus | TH |
| Hypothalamus | Hypothalamus | HY |
| Brainstem | Midbrain | MB |
|  | Pons | P |

#### Structural connectivity (SC) via fiber tractography

We used the same 21 nodes as in the rsFC analysis to assess SC between pairs of nodes using the streamlines fiber tractography pipeline described in [19, 33]. In brief, using the pre-processed data, we first estimated the fiber orientation distribution (FOD) map for each subject [39–41]. Then we considered a specific node as the seed and the remaining 20 nodes as individual target nodes and computed SC between each pair of nodes by seeding randomly within the seed mask (maximum 50,000 seeds - seeding termination criteria during streamline tractography) with a minimum streamline length of 3 mm, FOD cut-off 0.05, angle 45°, step size 0.025. This approach generated a 21 × 21 SC matrix. SC within the seed regions was ignored.

#### Statistical analysis

Statistical analyses were performed and visualized using MATLAB R2023a (www.mathworks.com) and GraphPad Prism version 10.2.2 for Windows (GraphPad Software, La Jolla, California, USA). Data were examined using a two-way ANOVA with strain and sex as fixed factors to assess the effects of strain, sex, and their interaction on FA, ALFF, rsFC, and SC, followed by Tukey’s or Holm-Šidák post-hoc analysis. Our primary dependent variables (i.e., regional FA, regional ALFF, SC, and FC) were examined for outliers defined as >2 standard deviation above or below the mean. We identified 2 C57-male, 1 DBA-male, and 1 DBA-female outliers, which were excluded from further analysis.

#### Correlation between rsfMRI and dMRI

Correlations between the brain structural and functional properties were examined using generalized linear regression analysis in MATLAB R2023a (www.mathworks.com) (Fig. 1d).

## Results

### Effects of strain and sex on brain amplitude of low frequency fluctuations (ALFFs)

First, we generated group-specific ALFF maps to observe changes in the ALFF patterns in each strain and sex (Fig. 2). At whole brain scale, C57 mice showed more pronounced ALFF (mean ± SD, female: 1.44 ± 0.13 and male: 1.58 ± 0.11) than DBA animals (female: 1.01 ± 0.08 and male: 1.01 ± 0.16). As the fMRI data were co-registered into the mouse brain atlas [19], next, we computed mean ALFF from 70 brain regions from each subject. Except agranular insular cortex (AI), where DBA mice showed greater activation than C57 mice, all other regions exhibited either higher ALFF in C57 or trivial changes between C57 and DBA mice. Finally, we quantified the effects of strain and sex on brain ALFF. We identified significant effects of strain or sex on ALFF in 12 brain regions out of 70 (2 × 2 ANOVA) as described below. In this analysis, no significant interactions were observed between strain and sex.

**Fig. 2:**
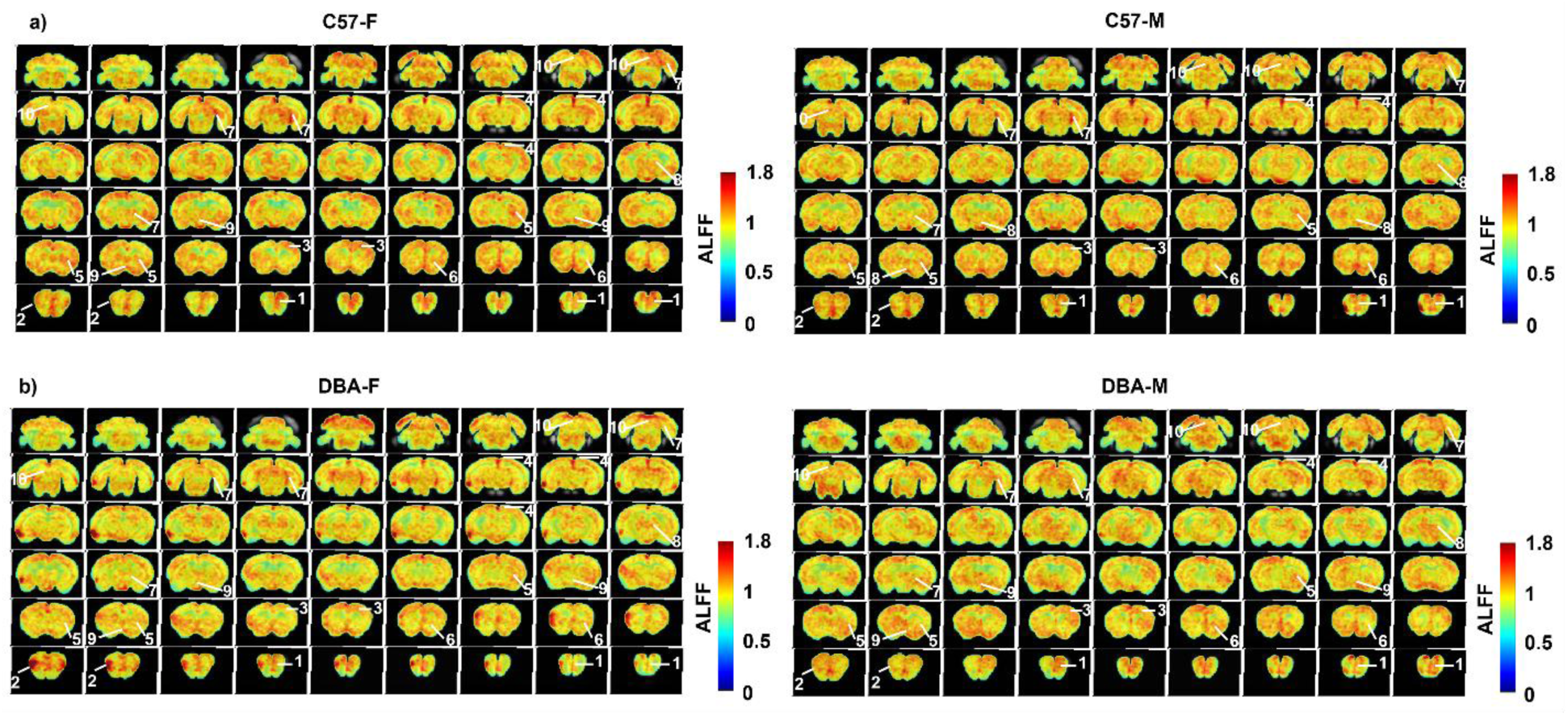
Average amplitude of low frequency fluctuation (ALFF) between male and female C57BL/6J and DBA/2J mice: a) C57BL/6J – female (left), and male (right). b) DBA/2J – female (left), and male (right). Hot and cold colors represent enhanced or suppressed ALFF, respectively. Anatomical regions were identified by mapping onto an in-house mouse brain atlas2^21^ (1: Prelimbic/ Infralimbic cortex (PL/ IL), 2: Agranular insular cortex (AI), 3: Motor/ Sensorimotor cortex (MO/ SS), 4: Caudate putamen (CP), 5: Nucleus accumbens (ACB), 6: Dentate gyrus (DG), 7: Thalamus (TH), 8: Hypothalamus (HY), 9: Midbrain (MB)).

In 8 brain regions, a significant main effect of strain, but not sex (p’s > 0.05), was observed (Fig. 3a, i-viii). C57 mice showed significantly enhanced ALFF compared to DBA mice in the prelimbic cortex (PL; F_1,30_ = 16.32, p = 0.0003), infralimbic cortex (IL; F_1,30_ = 11.91, p = 0.002), retrosplenial cortex (RSP; F_1,30_ = 5.27, p = 0.03), motor cortex (MO; F_1, 30_ = 17.77, p = 0.0003), somatosensory cortex (SS; F_1,30_ = 15.45, p = 0.0005), dentate gyrus (DG; F_1,30_ = 15.65, p = 0.0004), and midbrain (MB; F_1,30_ = 16.22, p = 0.0004). In only 1 brain region, the agranular insular cortex (AI) was ALFF significantly higher in DBA relative to C57 mice (F_1,30_ = 10.50, p = 0.003).

**Fig. 3:**
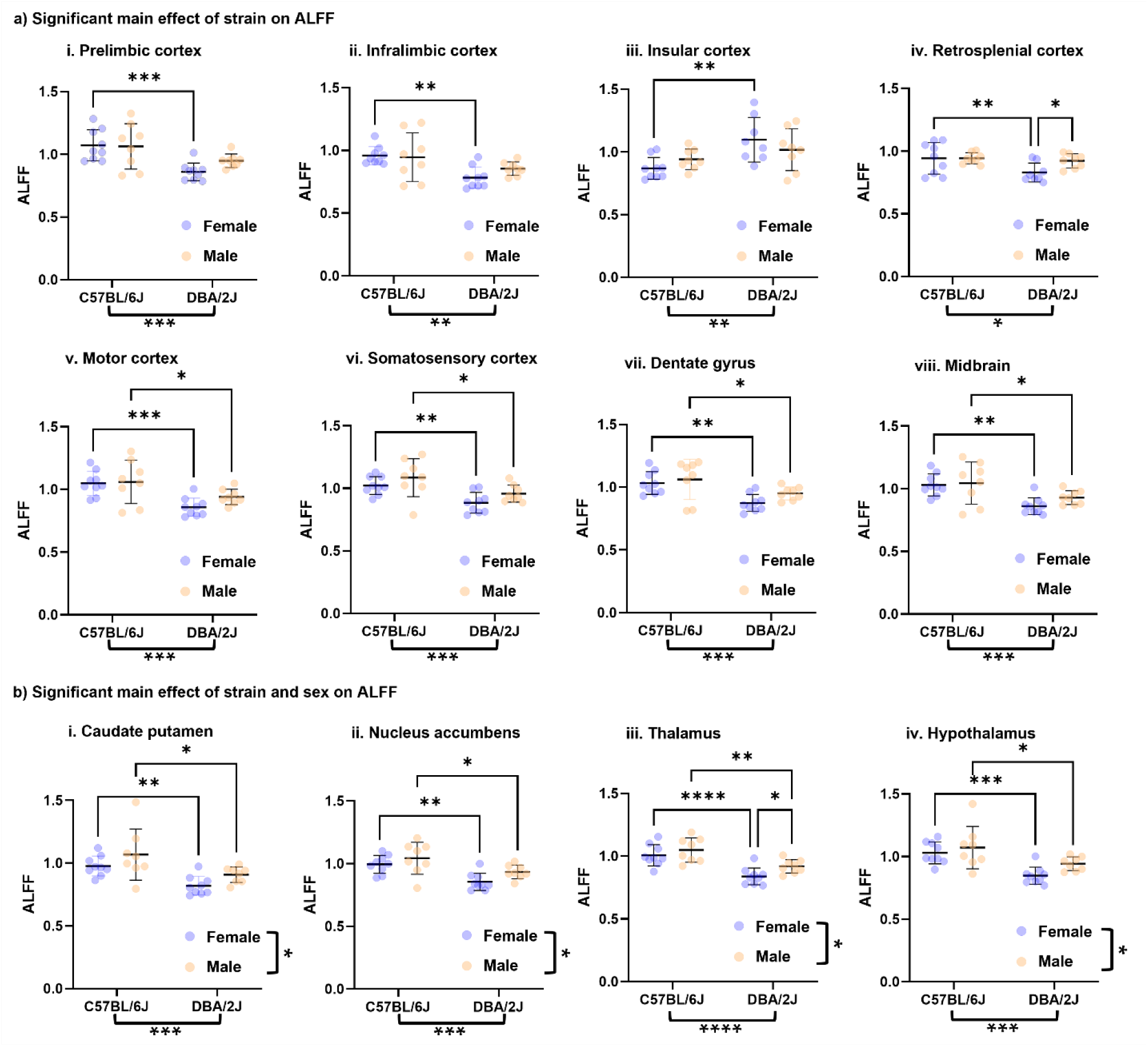
a) Significant main effect of strain on ALFF: prelimbic cortex (PL), infralimbic cortex (IL), insular cortex (AI), retrosplenial cortex (RSP), motor cortex (MO), somatosensory cortex (SS), dentate gyrus (DG), and midbrain (MB). b) Significant main effect of strain and sex on ALFF: caudate putamen (CP), nucleus accumbens (ACB), thalamus (TH), and hypothalamus (HY). Two-way ANOVA, α ≤ 0.05; *p≤0.05, **p<0.01, ***p<0.001, and ****p<0.0001.

In the remaining regions, both a significant main effect of strain and sex were observed (Fig. 3b, 1-iv). In the caudate putamen (CP; main effect of strain: F_1,30_ = 15.58, p = 0.0004; main effect of sex: F_1,30_ = 4.96, p = 0.03), nucleus accumbens (ACB; main effect of strain: F_1, 30_ = 18.47, p = 0.0002; main effect of sex F_1,30_ = 4.90, p = 0.03), hypothalamus (HY; main effect of strain: F_1,30_ = 19.13, p = 0.0001, main effect of sex F_1,30_ = 4.1, p = 0.05), and thalamus (TH; main effect of strain: F_1,30_ = 31.73, p < 0.0001; main effect of sex: F_1,30_ = 5.30, p = 0.03) C57 mice had greater ALFF relative to DBA animals and males had a more robust response compared to females.

### Effects of strain and sex on brain fractional anisotropy (FA)

Effects of strain and sex on brain FA were assessed using whole-brain voxel-based analysis (VBA). By comparing differences in voxel intensities across groups, we identified regional differences in the degree of water diffusion. A 2 × 2 ANOVA revealed a significant effect of strain on FA in numerous brain regions (minimum cluster size >25 voxels, FDR = 0.05, p < 0.006) (Fig. 4a). Decreased FA in DBA mice was most pronounced in several cortical regions, such as anterior cingulate cortex (ACA), MO, SS, RSP, visual (VIS), and entorhinal cortex (ENT). Similarly, DBA mice exhibited decreased FA in subcortical regions, including the basal ganglia (globus pallidus (GP) and CP, amygdala (AMY), and TH. Further, FA in the major white matter tracts, i.e., corpus callosum (cc) and internal capsule (int) of the DBA mice, was significantly reduced as compared to the C57. Regardless of the strain, a significant main effect of sex on FA (minimum cluster size >25 voxels, FDR = 0.05, p < 0.005) was observed in the subcortical brain regions and rostro-caudal cc (Fig. 4b). In males, FA was significantly reduced in the SS, TH, AMY, and cc (Fig. 4b – blue), whereas in females, frontal (olfactory (OLF), AI, piriform – PIR) and mid/hindbrain regions (superior colliculus – SC, midbrain reticular nucleus – MRN, pons – P) showed significantly reduced FA. In addition, we observed more pronounced strain differences in the males compared to females on FA, yielding significant strain × sex interactions in the prefrontal cortex (PFC), MO, SS, AI, RSP, ENT, CA1, AMY, HY, TH, and periaqueductal gray (PAG), as well as in the cc and int (Fig. 4c).

**Fig. 4:**
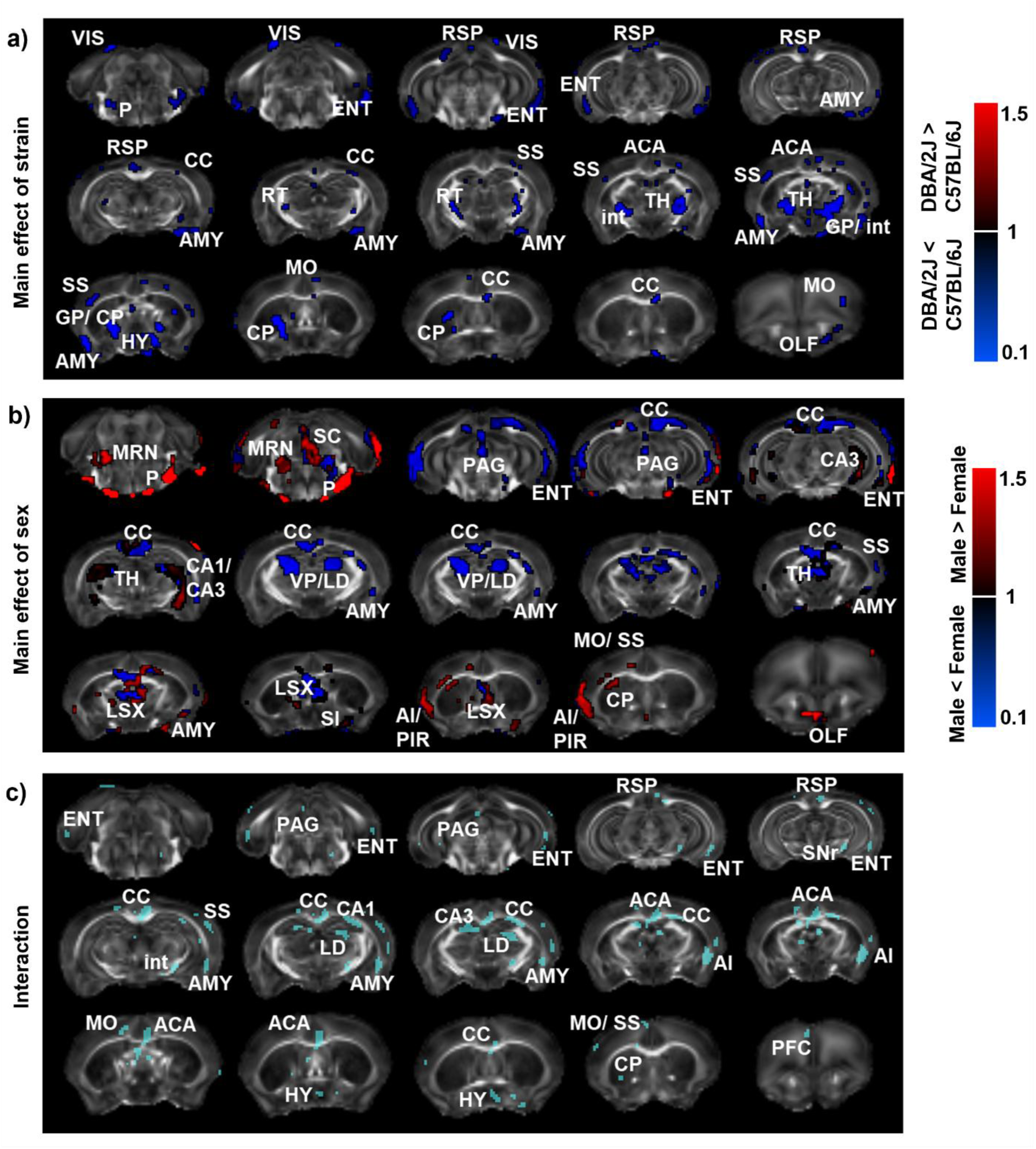
Strain and sex-dependent effects on fractional anisotropy (FA) in male and female DBA/2J and C57BL/6J mice. Data were analyzed using 2-factor ANOVA (α = 0.05, FDR corrected, n = 10 per strain and sex). a) Brain regions where a significant main effect of strain was observed: minimal cluster size >25 voxels, p<0.006. 1 = no change; <1 (blue) reduced FA in DBA/2J; >1 (red) increased in DBA/2J. b) Brain regions with a significant main effect of sex: minimal cluster size >25 voxels, p<0.05. 1 = no change; <1 (blue) reduced males >1 (red) increased in males. c) Interaction between strains and sex. c) Brain regions where there was an interaction between strain and sex on FA. ACA = anterior cingulate area, PL = prelimbic area, ILA = infralimbic area, PFC = prefrontal cortex, AI = agranular insular area, MO = motor area, SS = sensorimotor area, RSP = retrosplenial area, VIS = visual area, ENT = Entorhinal area, PIR = piriform area, CA1 = CA1 area of the hippocampus, CA2 = CA2 area of the hippocampus, CA3 = CA3 area of the hippocampus, DG = dentate gyrus, CP = caudate putamen, ACB = nucleus accumbens, LSX = lateral septal complex, GP = globus pallidus, AMY = amygdala, TH = thalamus, RT = Reticular nucleus of the TH, VP = ventral posterior complex of the TH, LD = Lateral dorsal nucleus of the TH, HY = hypothalamus, MB = midbrain, PAG = periaqueductal gray, SC = superior colliculus, MRN = midbrain reticular nucleus, SNr = substantia nigra, P = pons, CC = corpus callosum, int = internal capsule.

### Relationship between brain FA and ALFF

Next, we examined if spontaneous neural activity is governed by underlying microstructural integrity to gain insights into brain structure-function relationships. General linear regression analysis identified 6 regions from the cortical, limbic, and brainstem networks where ALFF and FA were significantly correlated in C57 and DBA mice. The identified areas include the RSP (R^2^ = 0.75, p < 0.0001) and MO (R^2^ = 0.56, p < 0.0001) from the cortex (Fig. 5a), MB (R^2^ = 0.15, p = 0.02) from the brainstem (Fig. 5b), and ACB (R^2^ = 0.71, p < 0.0001), AMY (R^2^ = 0.72, p < 0.0001), and TH (R^2^ = 0.74, p < 0.0001) from the limbic network (Fig. 5c). Positive correlation between FA and ALFF suggest that WM integrity is supportive to ALFF observed in these regions.

**Fig. 5:**
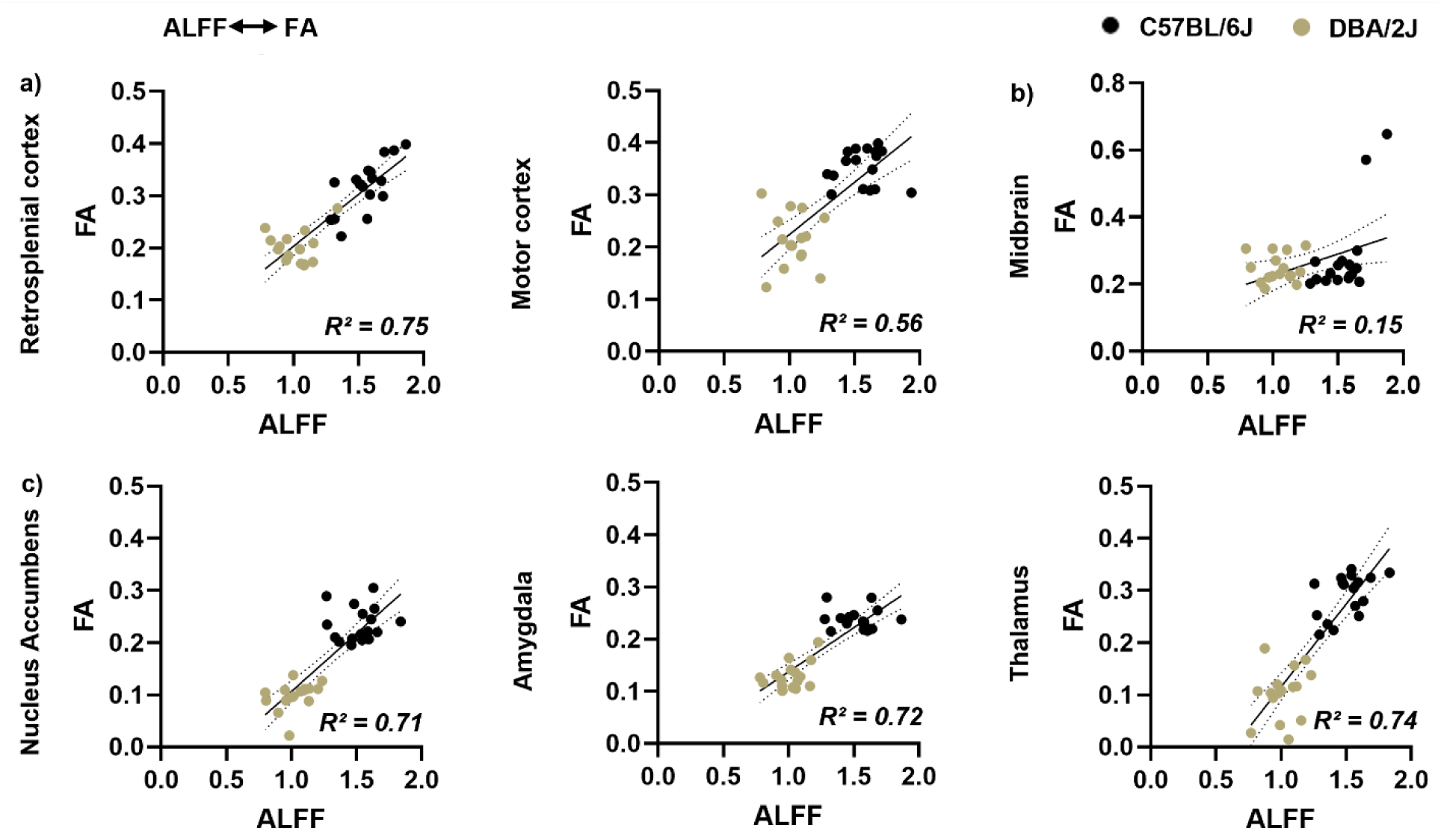
Brain regions where there was a significant correlation between the amplitude of low frequency fluctuation (ALFF) and fractional anisotropy (FA) (generalized linear regression, *p<0.05): a) cortical network: retrosplenial cortex and motor cortex, b) midbrain, and c) limbic network: nucleus accumbens, amygdala, and thalamus.

### Selection of nodes for functional and structural connectivity analysis

Based on regional brain properties (ALFF and FA), we identified 21 regions that exhibited significant differences in ALFF or FA properties between the strains or sexes (Table 1). These regions were then used as nodes to compute node-to-node FC and SC.

### Effects of strain and sex on rsFC

A representative image of the 21 brain regions used for connectivity analysis is shown in fig. 6a (black circles), and a representative connectivity network where nodes are interconnected via edges (black lines) is shown in fig. 6b. We observed a significant effect of strain (2 × 2 ANOVA) on several node-to-node rsFC as listed below (Fig. 6c and Table 2). Of the 14 identified pathways, the top 5 based on p-values were the SS - TH, AMY – ACB, SS – AMY, SS – RSP, and MO – AMY (Fig. 6e i-v). In all of these pathways, the strength of the connection was greater in C57 mice compared to DBA animals.

**Fig. 6:**
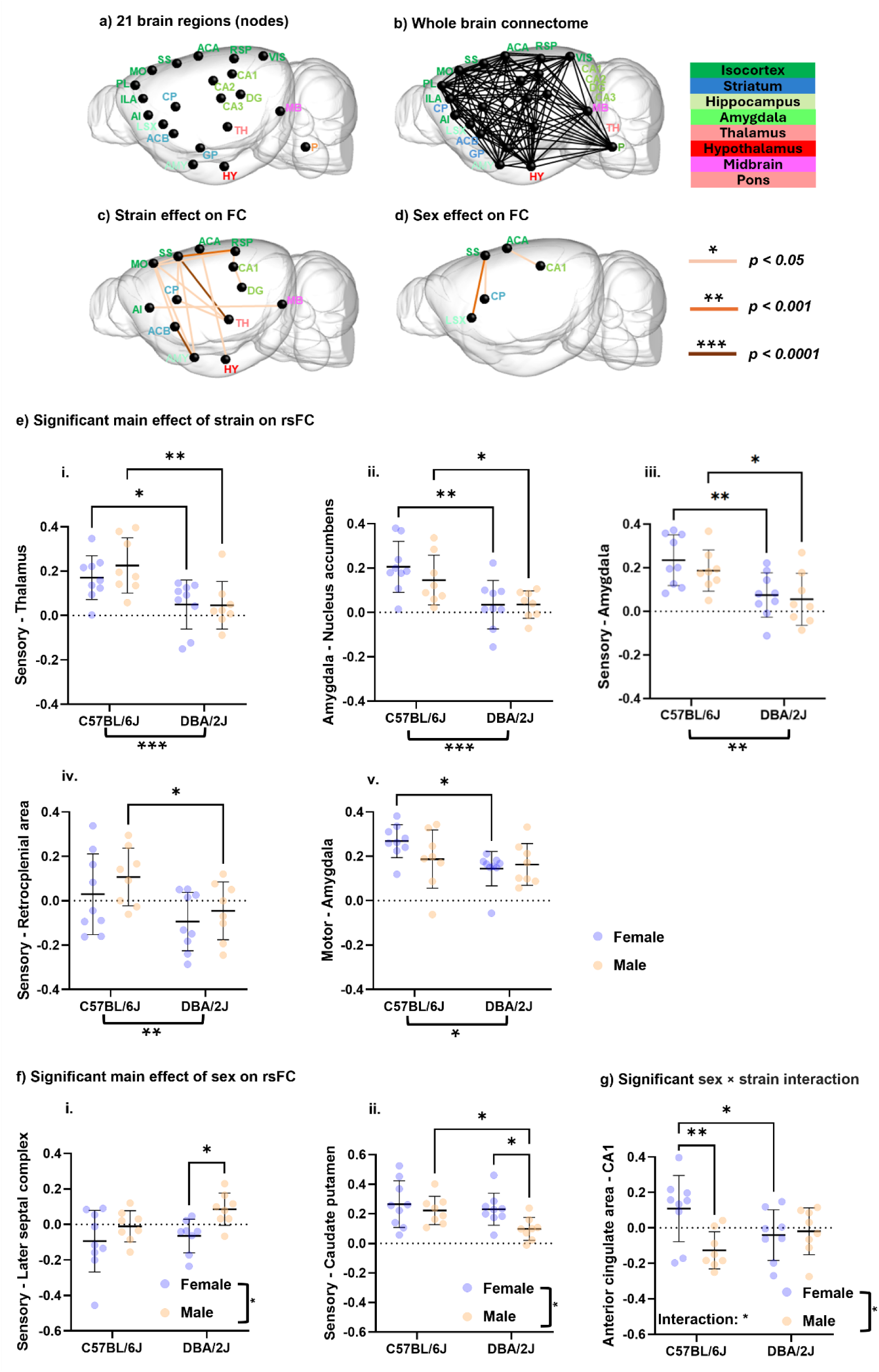
Functional connectivity analysis of male and female C57BL/6J and DBA/2J mice: a) 21 brain regions (black nodes) included in the node-to-node connectivity analysis, b) whole brain connectivity network among the 21 nodes. Black lines represent the connection between the nodes, c) functional connectivity (FC) between brain regions where a significant main effect of strain was observed, d) FC that was significantly altered by sex. For c-d, the color of the line represents the level of significance as denoted in the figure. Quantification of the significant effects on rsFC due to e) only strain but not sex, f) only sex but not strain, and g) significant sex × strain interaction. Statistical significance was tested via 2-way ANOVA, α ≤ 0.05. *p≤0.05, **p<0.01, ***p<0.001, and n.s.: not significant.

**Table 2:**
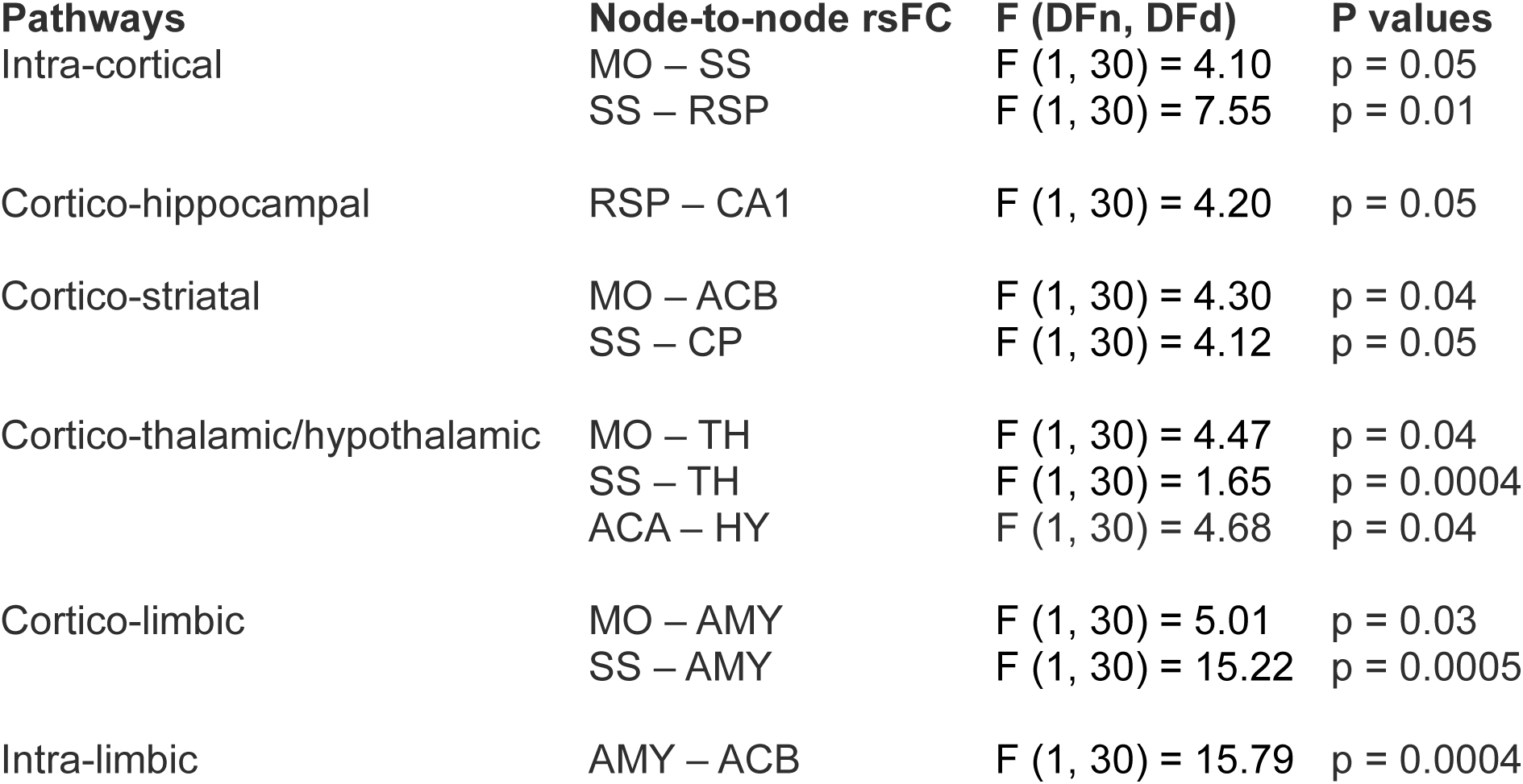
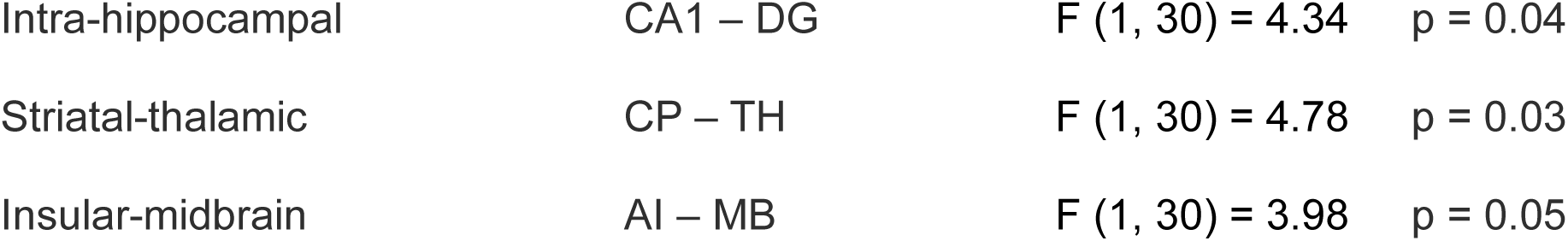
List of the rsFC between brain regions where a significant main effect of strain was observed.

| Pathways | Node-to-node rsFC | F (DFn, DFd) | P values |
| --- | --- | --- | --- |
| Intra-cortical | MO - SS | F (1, 30) = 4.10 | p = 0.05 |
|  | SS - RSP | F (1, 30) = 7.55 | p = 0.01 |
| Cortico-hippocampal | RSP - CA1 | F (1, 30) = 4.20 | p = 0.05 |
| Cortico-striatal | MO - ACB | F (1, 30) = 4.30 | p = 0.04 |
|  | SS - CP | F (1, 30) = 4.12 | p = 0.05 |
| Cortico-thalamic/hypothalamic | MO - TH | F (1, 30) = 4.47 | p = 0.04 |
|  | SS - TH | F (1, 30) = 1.65 | p = 0.0004 |
|  | ACA - HY | F (1, 30) = 4.68 | p = 0.04 |
| Cortico-limbic | MO - AMY | F (1, 30) = 5.01 | p = 0.03 |
|  | SS - AMY | F (1, 30) = 15.22 | p = 0.0005 |
| Intra-limbic | AMY - ACB | F (1, 30) = 15.79 | p = 0.0004 |
| Intra-hippocampal | CA1 – DG | $F(1, 30) = 4.34$ | $p = 0.04$ |
| Striatal-thalamic | CP – TH | $F(1, 30) = 4.78$ | $p = 0.03$ |
| Insular-midbrain | AI – MB | $F(1, 30) = 3.98$ | $p = 0.05$ |

Compared to the strain differences observed, the effects of sex on rsFC were subtle. Using 2 × 2 ANOVAs, significant main effects of sex, but not strain, were observed in two regions. In both the cortico-limbic (SS – lateral septal complex (LSX) and sensory-striatal (SS – CP) connections while males had a strong SS – LSX connection compared to females, the opposite was true in the SS – CP connection, where it was stronger among females relative to males (Fig. 6f i-ii).

Finally, in one connection, the cingulate-hippocampal ACA – CA1, there was a significant main effect of sex (F_1,30_ = 4.37, p = 0.04) and a significant sex × strain interaction (F_1,30_ = 6.32, p = 0.01) (Fig. 6g). Tukey’s post-hoc analysis showed a significant effect of strain in females (p = 0.04, Cohen’s d = 0.15), but not in males (p = 0.15, Cohen’s d = −0.11), and significant effect of sex within C57 mice (p =0.002, Cohen’s d = 0.24) (Fig. 6g).

### Effects of strain and sex on brain SC

We observed fewer strain effects on the brain-wide structural networks as compared to the rsfMRI networks. 2 × 2 ANOVA with Holm-Šidák post-hoc analysis revealed significant strain and sex effects predominantly in the cortico-limbic, cortico-thalamic, and insular-midbrain networks (Fig. 7 a,b). Structural connections between each pair of nodes represent the streamlines connecting the respective nodes derived via fiber tractography and were overlaid on a 3D glass brain (Fig. 7 c-f i). Red, green, and blue colors represent the direction of the fibers in x, y, and z axis.

**Fig. 7:**
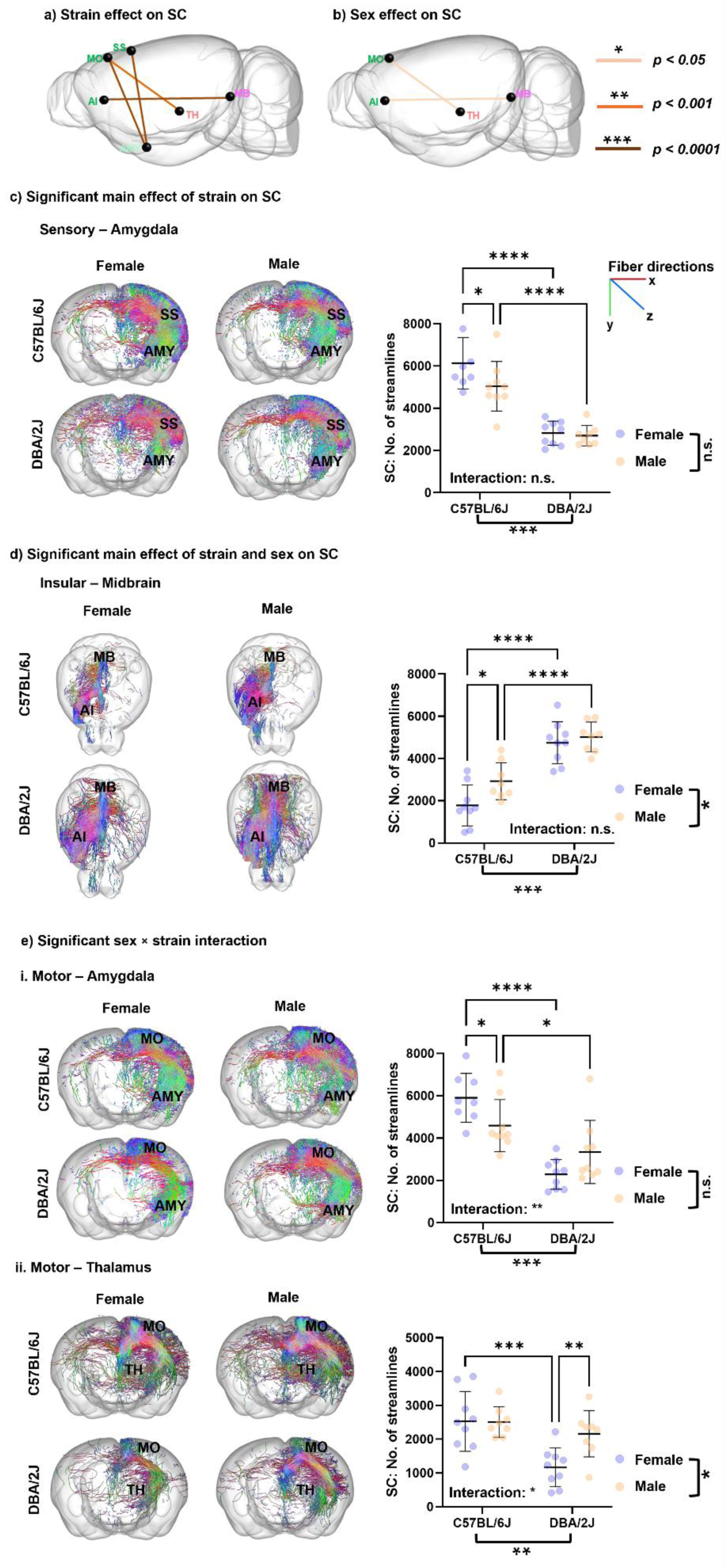
Structural connectivity analysis of male and female C57BL/6J and DBA/2J mice: a) Structural connectivity (SC) between brain regions where a significant main effect of strain was observed, and b) SC that was significantly altered by sex. For a-b, the color of the line represents the level of significance as denoted in the figure. Tractography maps and the number of streamlines connecting two specific nodes in male and female C57BL/6J and DBA/2J mice: c) significant main effect of strain on SS – AMY SC, d) significant main effect of strain and sex but not interaction on AI – MB, e) significant sex × strain interaction on i. MO-AMY, and ii. MO-TH SC. Red, green, and blue colors in the tractography maps represent fibers oriented in x, y, and z directions. Statistical significance was tested via 2-way ANOVA, α ≤ 0.05. *p≤0.05, **p<0.01, ***p<0.001, and n.s.: not significant.

The SS–AMY pathway was significantly altered by strain (F_1,30_ = 79.55, P < 0.0001) but not by sex or interaction. C57 mice had significantly more streamlines in this network compared to DBAs (Fig. 7c). In the salience-midbrain network (AI – MB) there was a significant effect of strain (F (1, 30) = 66.81, P < 0.0001) and sex (F (1, 30) = 5.30, P = 0.02) but the interaction between these factors was not significant (Fig. 7d). Streamlines were greater in male animals relative to females, and in DBA mice relative to C57s.

In 2 structural connections, where a significant interaction between strain and sex was observed. In the cortico-limbic (MO-AMY) connection, there was a significant main effect of strain (F (1, 30) = 37.16, p < 0.0001) and interaction between sex and strain (F (1, 30) = 8.80, p = 0.006) (Fig. 7e-i). Holm-Šidák post-hoc analysis revealed a significant effect of strain in females (p < 0.0001, Cohen’s d = 3620) and males (p = 0.03, Cohen’s d = 1250), as well as sex within the C57 group (p = 0.02, Cohen’s d = 1310) (Fig. 7e-i). Additionally, in a cortico-thalamic connection (MO – TH), significant main effects of strain (F (1, 30) = 13.67, p = 0.001), sex (F (1, 30) = 4.42, p = 0.04), and their interaction sex and strain (F (1, 30) = 4.82, p = 0.03) were detected (Fig. 7e-ii). Tukey’s post-hoc analysis showed a significant effect of strain in females (p = 0.0002, Cohen’s d = 1363) but not in males (p = 0.31, Cohen’s d = 347.3) and a significant effect of sex within the DBA group (p < 0.005, Cohen’s d = −994.1) (Fig. 7e-ii).

### Relationship between brain SC and rsFC

Next, we tested whether the observed changes in the FC networks could be seen in the underlying SC networks using streamlines fiber tractography. Using generalized linear regression analysis, we identified four pathways, including the MO – AMY (R^2^ = 0.15, p = 0.02), SS – AMY (R^2^ = 0.72, p < 0.0001), MO – TH (R^2^ = 0.15, p = 0.02), and AI – MB (R^2^ = 0.72, p < 0.0001) that show significant correlations between rsFC and SC (Fig. 8 a-d).

**Fig. 8:**
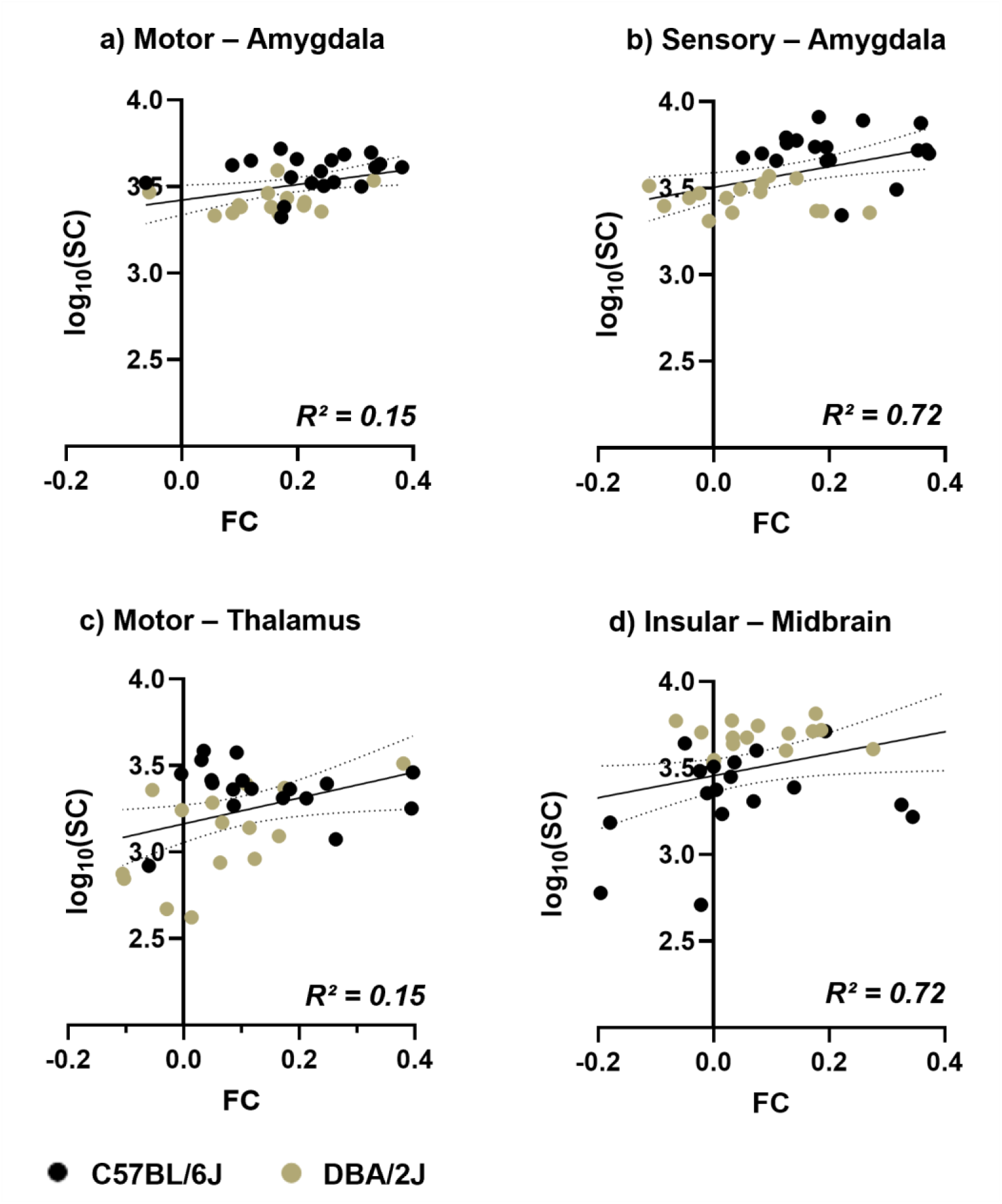
Correlation between the FC and SC (generalized linear regression, *p<0.05) of male and female C57BL/6J and DBA/2J mice: a) motor – amygdala, b) sensory – amygdala, c) motor – thalamus, and b) insular – midbrain.

## Discussion

In this study, we conducted a systematic examination of the functional and structural architectures of the two most widely used mouse strains, C57 and DBA, in neuroscience research. We detected significant differences between strains and sexes in ALFF and FA in several brain regions. These include the prefrontal, sensory-motor, default mode network (DMN), salience, visual, hippocampal, basal ganglia, limbic, and midbrain/hindbrain. From connectivity analysis, we further identified significant effects of strain and sex on node-to-node FC and SC within brain pathways, including cortico-thalamic, cortico-limbic, and salience-midbrain. Importantly, observed changes in brain functional properties (ALFF and FC) were governed by the changes in the microstructural fingerprints (FA and SC). Together, this study identifies baseline differences in the brain connectome specific to strain and sex that should be considered while interpreting the behavioral differences between these strains of mice.

rsfMRI-based ALFF is a fundamental measure of the brain that reflects spontaneous neural activity at resting state. These frequency-dependent alterations can vary between brain regions or subjects and can serve as a marker of individual differences or pathology [42, 43]. dMRI-based FA, on the other hand, measures the overall directionality of water diffusion that reflects white matter integrity in the brain. Deviations in FA values indicate damage in the myelin sheath or changes in the intra-voxel crossing fibers, fiber density, or diameter, resulting in changes in the structural fingerprints underlying brain networks [44, 45]. Therefore, both ALFF and FA play substantial roles in shaping brain connectomics, which defines the spatial connectivity patterns of neural pathways. In this study, we utilized ALFF, FA, FC, and SC as markers to capture strain- and sex-specific connectome signatures of C57 and DBA mice by combining non-invasive rsfMRI and dMRI techniques.

In the 1950s, Rodgers and McClearn were some of the first to demonstrate behavioral differences between C57 and DBA mice. In this study, it was demonstrated that C57 mice showed a preference for an alcohol-containing solution, while DBA mice avoided it[46]. Since this time, divergent traits for these strains have been repeatedly observed for phenotypes related to addiction [15, 17, 47], learning and memory[12, 13], and emotionality [48, 49], among others. Notably, some of these phenotypic differences are observed immediately and without manipulation, suggesting that innate differences between the strains drive these effects. These strains and derivatives resulting from crosses between them have been widely used to demonstrate that genetic differences underlie these behavioral traits.

Identified genetic differences between these strains likely influence the brain function that drives the observed behavioral differences. Both structural and functional differences in the brain have been observed between C57 and DBA mice. As one example, strain differences in hippocampal function have been reported [50]. Differences in brain structure have also been identified between these strains. Prior research has demonstrated reduced cc size in DBA mice relative to C57s [51, 52]. The current work verifies this difference, which was initially observed and identified using histological methods. Importantly, though, the use of MRI by our group and others allows for a brain-wide analysis [24]. In this manuscript, we extend prior research and observe *in vivo* functional differences across the brain between these strains. We also demonstrate that some of the functional changes identified correlate with structural changes, providing one mechanism by which genetic differences between these strains can alter behavioral outcomes.

Our results on strain differences in baseline measurements of functional and structural brain properties are consistent with research in humans, indicating a genetic component to these traits. For example, the heritability of functional connectivity in the DMN in humans has been estimated to be approximately 0.5 [53, 54]. Moreover, several groups have identified a genetic contribution to the structure of cortical and subcortical brain regions [55, 56]. In parallel to this human research, work from Wang and colleagues have detailed the structure of 4 inbred mouse strains, including C57 and DBA, using DTI [24]. Notably, sevral key findings were reproducible in our in-vivo study, including weaker FA in the CC, ACA, and SS of DBA than C57, as well as trivial effect of sex on diffusion metrics compared to the strain differences. While this work offers a greater spatial resolution than current methods, it requires approximately 23 hours of MRI scan time per sample and, importantly, must be performed *ex vivo*. In the current work, we utilized an *in vivo* imaging protocol that allow us to measure both the structural and functional properties of the brain during the same imaging session and within the same animal. Further, our MRI scan was completed in an hour, allowing for a high throughput.

By using this *in vivo* imaging approach, we can explore the relationship between structural and functional properties of the brain within the same individual. Here, we saw a positive relationship between brain activity as measured by ALFF and white matter structural integrity as measured by FA in 6 brain regions. In humans, research has identified the DMN as an interconnected network in the brain that is active when the subject is at rest and not performing a task [57]. Researchers have identified a similar network in the rodent brain that is identified by several cortical areas, including the RSP [58]. The RSP in rodents is similar to the posterior cingulate cortex in humans and a primary hub of the DMN. In our work, we see activation of the RSP. Moreover, we identified a strong correlation between ALFF and FA within the RSP, a finding that replicates human imaging research [59]. The DMN has also been highly studied as a potential network that drives human behaviors. As examples, this network has been linked to anxiety, memory, cognition, and addiction [60–62], many of which are divergent between C57 and DBA mice.

While we observed some sex differences in this study, these were smaller in magnitude than the observed strain differences. These findings are in line with prior research that found more strain differences in structural brain measures between strains than sex differences [24]. Additionally, research examining behavioral outcomes has similarly observed more differences between strains than between sexes [63] within a strain. While we identified relatively few sex differences in brain function or structure, it is important to note that we detected sex differences in brain regions that have been observed in humans. For example, although there are inconsistencies in the field, there have been several studies that observed the cc to be larger in females compared to males [64]. In one study, this difference was observed in children as young as one month of age after controlling for brain volume [65]. Thus, these findings are translationally relevant to humans.

### Conclusion

In this study, we utilized *in vivo* MRI to examine structural and functional differences between awake male and female C57 and DBA mice. We observed differences in these parameters between strains and sexes. Importantly, these data replicate findings from human neuroimaging studies, indicating the translational value of this work. Additionally, they provide a brain-wide view of baseline differences between two of the most widely used strains in behavioral genetics research. Utilizing this high-throughput MRI protocol, which can be combined with behavioral testing or disease models, may provide insight into brain mechanisms.

## Data Availability Statement

Data are available from the authors by request.

## Acknowledgments

This work was supported by the National Institute on Drug Abuse (DA060335, H.M.K.), Penn State’s Department of Biobehavioral Health, Social Science Research Institute, and Consortium on Substance Use and Addiction. The authors would like to acknowledge the Huck Institutes High Field Magnetic Resonance Imaging Core Facility (RRID:SCR_024461) for use of their Bruker Biospec 70/30. The content of this manuscript is solely the responsibility of the authors and does not necessarily represent the official views of the National Institute on Drug Abuse or the National Institutes of Health.

